# Phage Steering in the Presence of a Competing Bacterial Pathogen

**DOI:** 10.1101/2024.09.28.615603

**Authors:** Sean Czerwinski, James Gurney

## Abstract

The rise of antibiotic-resistant bacteria has necessitated the development of alternative therapeutic strategies such as bacteriophage therapy, where viruses infect bacteria, reducing bacterial burden. However, rapid bacterial resistance to phage treatment remains a critical challenge, potentially leading to failure. Phage Steering, which leverages the evolutionary dynamics between phage and bacteria, offers a novel solution by driving bacteria to evolve away from virulence factors or resistance mechanisms. In this study, we examined whether Phage Steering using bacteriophage Luz19 could function in the presence of a competing pathogen, *Staphylococcus aureus* (USA300), while targeting *Pseudomonas aeruginosa* (PAO1). Through *in vitro* co-evolution experiments with and without the competitor, we observed that Luz19 consistently steered *P. aeruginosa* away from the type IV pilus (T4P), a key virulence factor, without interference from *S. aureus*. Genomic analyses revealed mutations in T4P-associated genes, including *pilR* and *pilZ*, which conferred phage resistance. Our findings suggest that Phage Steering remains effective even in polymicrobial environments, providing a promising avenue for enhancing bacteriophage therapy efficacy in complex infections.

**Importance:** Phage Steering—using phages that bind essential virulence or resistance-associated structures— offers a promising solution by selecting for resistance mutations that attenuate pathogenic traits. However, it remains unclear whether this strategy remains effective in polymicrobial contexts, where interspecies interactions may alter selective pressures. Here, we demonstrate that *Pseudomonas aeruginosa* evolves phage resistance via loss-of-function mutations in type IV pilus (T4P) when challenged with the T4P-binding phage Luz19, and that this evolutionary trajectory is preserved even in the presence of a competing pathogen, *Staphylococcus aureus*. Phage resistance was phenotypically confirmed via twitching motility assays and genotypically via whole genome sequencing. These findings support the robustness of Phage Steering under interspecies competition, underscoring its translational potential for managing complex infections—such as those seen in cystic fibrosis—where microbial diversity is the norm.

## Introduction

The increasing prevalence of antibiotic-resistant bacteria necessitates the exploration of alternative therapeutic approaches. Bacterio(phage) therapy, which utilizes viruses that infect and kill bacteria offers a promising avenue. However, the rapid emergence of phage-resistance *in vitro* raises alarms over phage therapy’s long-term efficacy [1, 2]. The antagonistic co-evolutionary dynamics of bacteria-phage interactions generates diversity in both the bacterial and phage populations, with reciprocal evolution of defense vs attack alleles occurring in an almost turn-based competition known as Arms Race Dynamics (ARD) [3-5] wherein bacterial resistance to phage emerges, followed by the evolution of phage infectivity [6-11]. Ultimately, in the lab, ARD culminates in the emergence of a resistant genotype that the phages cannot overcome [8], in most cases coming in the form of *de novo* mutations resulting in the loss or structural and functional attenuation of the specific surface receptors to which the bacteriophage binds [8]. Co-evolutionary dynamics between phage and bacteria can also progress from ARD to exhibit fluctuating selection dynamics in which bacteria resistance is not generalized [12]. The shortcoming of bacteriophages is to be expected, as Lenski and Levin proposed, there is an asymmetry in the evolutionary potential between the bacterium and its phage [8, 9]. This asymmetry is due, in part, to a single spontaneous *de novo* mutation resulting in significant receptor structure alteration and therefore acquisition of resistance for the bacterium, whereas the bacteriophage would need multiple successive *de novo* mutations in order to overcome resistance [13, 14]. Phage Steering seeks to exploit the selection of the host in response to phage pressure. In the event that resistance evolves against the phage, the *de novo* mutations that result should lead to the loss or functional attenuation of the binding receptor [9, 15, 16]. Phage Steering utilizes bacteriophages with known receptors, or factors, that are implicated in antimicrobial resistance or in virulence [15].

The intricate dynamics of bacteria-phage interactions, particularly the rapid emergence of phage resistance, pose significant challenges to the long-term efficacy of bacteriophage therapy. This is especially critical when considering pathogens like *Pseudomonas aeruginosa* (*PA*), a notorious Gram-negative bacterium known for its intrinsic multi-drug resistance and its role in chronic, often life-threatening infections [17]. Furthermore, *PA* infections often occur in a poly-microbial context, most notably alongside *Staphylococcus aureus* (*SA*), significantly complicating treatment strategies. These two pathogens are the most prevalent in cystic fibrosis infections, with studies indicating enhanced virulence when they coexist [18, 19]. The dynamic and often synergistic interactions between *PA* and *SA* in co-infection models [18, 20-23], emphasize the complexity of *PA*-associated polymicrobial infections. The community interactions found within a polymicrobial infection has the potential to greatly impact phage therapies and steering [24, 25]. *PA* is an opportunistic pathogen, and likely did not evolve its vast armament of virulence factors and antimicrobial resistance mechanisms in order to colonize the CF lung, but rather to outcompete other micro-organisms in the environment [26-29]. Thus, the question remains whether Phage Steering will prove efficacious in the presence of competing co-infecting organisms that exert their own selective pressure via competition or otherwise interaction.

In this study, we focus on the interplay between *Pseudomonas aeruginosa* - PAO1 (*PA)* and *Staphylococcus aureus* - USA300 (*SA*) and the phage Luz19. We employ the phage, Luz19, to steer PAO1 away from the expression or function of the phage receptor, the type IV pilus (T4P) [30, 31]. Co-evolving with and without the phage and *SA* present we determined if Luz19 is capable of performing Phage Steering on *PA* during bacterial competition. Our phenotypic assays of T4P activity and whole genome sequencing of evolved lines show that phage resistance was mediated by mutations in the Type IV pili system implying Phage Steering still works under competition with *SA* in LB.

## Materials and Methods

### Bacterial culture and phage preparation

*Pseudomonas aeruginosa* (Nottingham *PA*O1) and *SA* (USA300) were independently grown at 37°C with agitation (shaking at 200 rpm) in Lysogeny Broth (LB) media (25g^-L^ LB Broth (Lennox)). For phage stock quantification, Luz19 phage was diluted in 1X Phosphate Buffer Saline (PBS) and spot plated onto *PA*O1 soft lawns on Lysogeny Broth Agar (LBA, Agar 15g^-L^) and incubated at 37°C overnight. Plaque Forming Units (PFUs) were counted, and PFU/mL was calculated. Phage stock was diluted in 1X PBS. The final concentration of phage stock prior to initial inoculation was approximately 3.0e9 PFU/mL. Phage inoculation dosage was calculated for a multiplicity of infection (MOI) of 3, from estimated Colony Forming Unit (CFU) per mL by multiplying the Optical Density at a wavelength of 600 nm OD_600_ by 8.0e8.

### *In vitro* evolution experiment

We investigated bacterial evolutionary response to an anti-virulence bacteriophage, Luz19, over the course of 10 serial transfers (10 days) in the context of presence or absence of a common competitor, *SA*. A total of three co-evolution conditions were used, *PA* with *SA, PA* with Luz19 phage, and *PA* with both phage and *SA*. A control for serial transfer culturing was included with only *PA* serially transferred for the total 10 transfers. All co-evolution cultures (5 replicates) were conducted in 1 ml LB in 24 well plates (Corning)®) and incubated at 37°C, shaking, for 24 hours. Bacteria colony forming unit (CFU) estimates for initial and subsequent (for *SA*) inoculations were made using OD_600_ measurements. For accurate CFU estimation from OD_600_, *PA*O1 and USA300 suspended cultures were spectrophotometrically measured and then serially diluted and CFU plate counted. An OD_600_=1.0 was found to contain approximately 8.0e8 and 1.0e8 CFU/mL for *PA*O1 and USA300 respectively. 50 µL of *PA*O1 (1.6e7 CFU) was inoculated into wells containing 900 µL of LB media. In conditions with phage, bacteria were exposed to the 16 µL of the initial ancestral Luz19 phage at an MOI of 3:1 (4.9e7 PFU); from Luz19 stock at 3.0e9 PFU/mL at time point 0 (T0).

In conditions with competing *SA*, 50 µL of USA300 (2.0e6 CFU) was inoculated into the wells containing 900 µL of LB media and 50 µL of newly inoculated PAO1. In +*SA* conditions, 50 µL of *SA*, 5.0e6 CFUs, was re-introduced at each transfer to maintain the selective pressure generated by consistent competition.

In pilot experiments, we found that *SA* went extinct after no more than 3 transfers when co-cultured with PAO1 at comparable relative concentrations (data not shown). After 24 hours, 100 µL was transferred into 900 µL of LB media. 300 µL was stored in 25% final concentration glycerol at – 80°C.

In the case that a replicate culture needed to be restarted, the frozen stock was used to resume the replicate from the most recently stored time point. Due to human error during transferring the *PA* (-/-) T5 replicates C, D, E were restarted from T4 for each respective replicates. Likewise, *PA* + Luz19 T8 E, restarted from T7 for respective replicate. 1 µl of Frozen culture was inoculated into 1 mL of LB and incubated at 37°C with shaking for 4 hours. 100 µL was then transferred into 900 µL of LB and normal culturing conditions resumed. Upon culmination of co-evolution culturing, all Transfer 10 replicates were frozen in glycerol (25% final) and stored at -80°C.

### Preparation for Twitching Motility and Growth Assays

Frozen replicates were cultured in LB at 37°C shaking overnight. The overnight culture was then transferred into a 1.5 mL microcentrifuge tube, briefly vortexed to break up aggregates, and then centrifuged at 17,000 RCF for 2 minutes to pellet the cells, then washed 2 times in 1X PBS and re-pelleted. Cells were resuspended in 1 mL of LB and further diluted in LB to an OD_600_ of 0.05. In the case of USA300-positive co-cultures, overnight cultures were streaked onto Pseudomonas Isolation Agar (PIA) and incubated at 37°C overnight to remove *SA*. Single colonies of isolated *PA* were incubated overnight at 37°C in LB.

### qPCR of Phages

qPCR of phage stock was performed using the QuantStudio 3 rt PCR system and analyzed with the ThermoFisher applied biosystems app. See SI for list of primers used.

### Twitching Motility Assays

We performed a twitching motility assay adapted from Alm & Mattick’s 1996 protocol, with the following modifications [32]. A p20 pipette tip was introduced to the cultures then pushed through a LB agar plate until the pipette tip contacted plastic at the base of the plate and then promptly removed. Plates were incubated at 37°C for 20 h. Agar was removed to reveal the imprint of *PA* growth. Imprints were stained with Crystal Violet for 15 min. Excess stain was gently washed off using sterile diH_2_O. Plates were then air dried. Once dry, the stained imprints were measured across at the widest point. All phenotypic assays were done in triplicate for each co-evolution replicate for a total of 15 assay replicates-per-condition.

Data transformations (log and Box-Cox using the car package for R 4.2.2 [33]) were attempted, but normality was not achieved across all treatments according to a Shapiro-Wilk test. Due to heteroskedasticity we used the Kruskal-Wallis test. Pseudo replication was corrected by first averaging the measurements taken of triplicates. When significant differences were found, post-hoc analyses were performed using the Dunn’s test with Bonferroni correction for Kruskal-Wallis. All statistical analysis was performed in R 4.2.2.

### Growth Assays

We performed a series of growth assays for *PA* of the four evolution conditions and the ancestor. All lines of the conditions were prepared following established protocols. They were then inoculated into 180 µL of LB in individual wells of a 96 well plate (VWR®), at an initial concentration of approximately 8.9e4 CFU per well. Each of the five lines per condition were subject to four treatments for the growth assays (All Condition x Line x Treatment combinations were performed in triplicate): no additional treatment (control), *SA* added at a 1:1 CFU ratio to *PA* (8.9e4 CFU), ancestral Luz19 phage added at an MOI of 3 (2.7e5 PFU), or both *SA* and Luz19 added. After inoculation with condition lines and treatments, the 96-well plates were loaded into an Agilent BioTek BioSpa^™^ Automated Incubator. The plates were incubated statically at 37°C to promote *PA* growth under the controlled treatment conditions. To monitor bacterial growth, the plates were automatically transferred every 30 minutes into an Agilent BioTek Cytation^™^ for OD_600_ measurements. A total of 49 readings per well were collected over a period of 24.5 hours. The temporal OD_600_ data were subsequently used to analyze growth dynamics under each treatment condition.

### Growth Analysis

Analysis of growth assays was performed using the Growthcurver package for R [34]. Growthcurver fits the data points collected for each individual replicate’s growth curve to a standard form of the logistics equation used in ecology and evolution [34]. Growth curves had the area under the growth curves differentiated for each co-evolution condition subject to four different growth assay treatments, with standard error bars. AUC is the area under the curve, calculated by summing the areas beneath the experimental curve obtained from the input measurements using the Growthcurver package [34].

The growth curves of *PA* from different co-evolution conditions under various treatments was assessed using the area under the curve (AUC). Statistical analyses were performed to determine significant differences in growth between conditions and treatments. Data transformations (log and Box-Cox using ‘car’ and ‘MASS’ packages for R 4.4.1 respectively[33]) were attempted but did not achieve normality across all treatments. We corrected for pseudo replication by averaging every three technical replicates within each Treatment x Condition to yield independent pseudo replicate values for AUC. To account for the non-normality observed in our data, we used the ARTool package for R 4.4.1 to perform a non-parametric two-way analysis of variance with Treatment and Condition as fixed factors. Subsequent post-hoc analysis was conducted within the ARTool framework to generate pairwise comparisons (with Bonferroni correction) for the Treatment x Condition interaction. No differences were observed in the control evolution line (**SI Fig.1**). Similar analyses were performed on intrinsic mean growth rate (r) and carrying capacity (k). (**SI Fig. 1 & 2**).

### DNA Extraction & Purification

Single colonies, one from each replicate line, were cultured in LB shaking overnight at 37°C. DNA extraction, isolation, and purification was performed using the Promega Wizard ® Genomic DNA Purification Kit following the manufacturer’s instructions.

### Sequencing

Whole genome sequencing of single colonies from each replicate was performed by SeqCoast Genomics (Portsmouth, NH, USA). SeqCoast prepared samples using the Illumina DNA Prep tagmentation kit and unique dual indexes. Sequencing was performed on the Illumina NextSeq2000 platform using a 300-cycle flow cell kit to produce a 2×150bp paired reads. 1-2% PhiX control was spiked into the run to support optimal base calling. Read demultiplexing, read trimming, and run analytics were performed using DRAGEN v4.2.7, an on-board analysis software on the NextSeq2000.

### Genomic Analysis

Genomic analyses were completed using breseq 0.38.3 [35] computational pipeline to identify mutations relative to the reference genome Pseudomonas aeruginosa PAO1 [36]. Sequencing of our ancestral nPAO1 genome and subsequent breseq analysis was used to identify single nucleotide polymorphisms (SNPs) that are not present in the reference genome [36]. We compared the ancestral nPAO1 and the transfer control (*PA* t10) to the evolved *PA* to remove SNPs that were a result of adaptation to the media.

## Results

### Twitching Motility

Selection by Luz19 phage attenuates twitching motility function in the presence of competing *SA* (**Fig. 1**). Twitching motility was attenuated in phage-positive conditions in the presence and absence of competing *SA* (p < 0.05 for both conditions), relative to the ancestral PAO1. There was no significant difference between the two positive-phage conditions nor between the ancestor and the negative/negative culture control. *PA* that was co-cultured with *SA* (*PA+SA*) did not have a significant difference in twitching motility relative to the ancestral PAO1 (p > 0.05).

**Fig 1.**
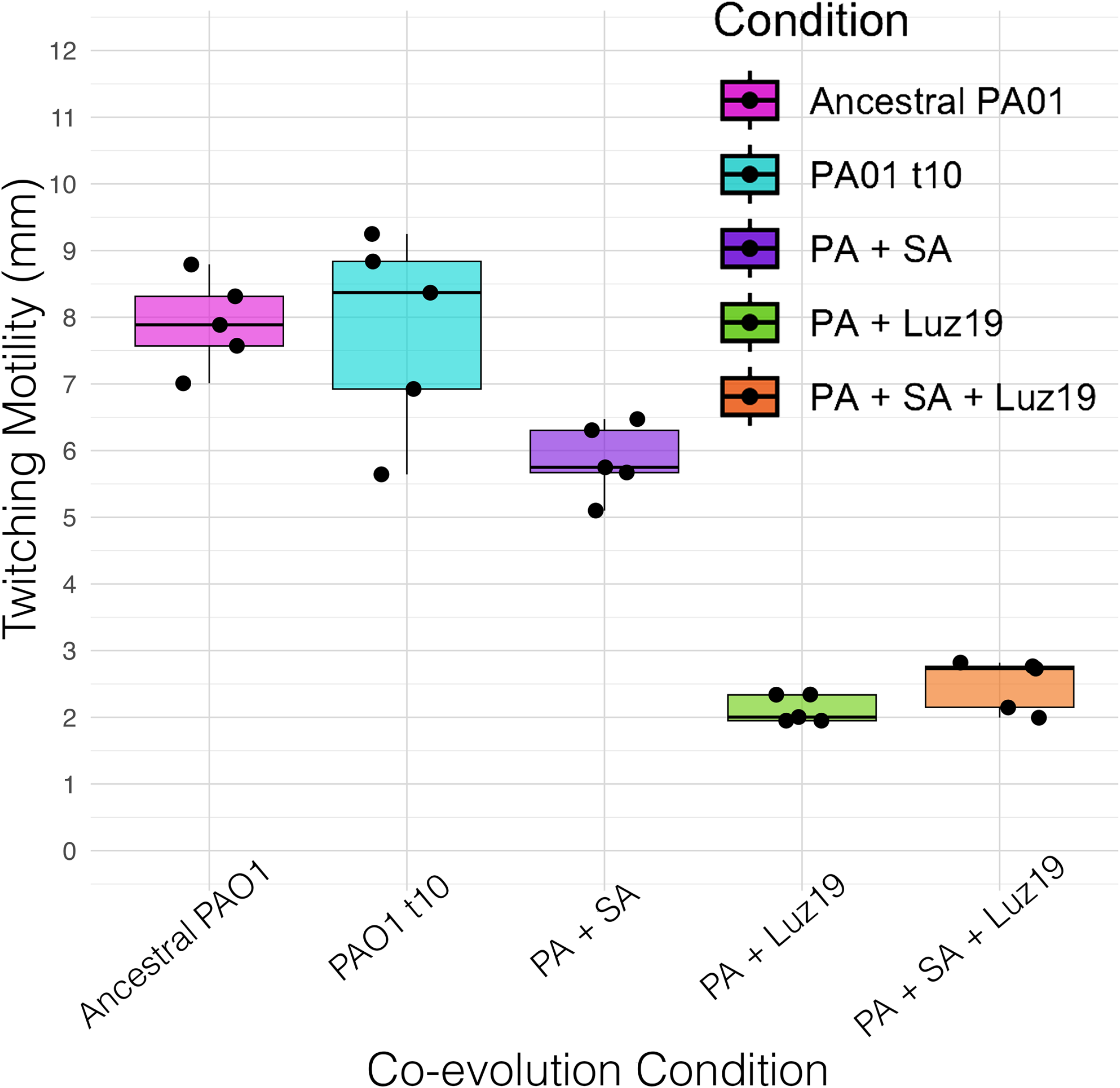
Twitching motility compared across evolution conditions. *PA* twitching motility decreased significantly (p < 0.05) in conditions exposed to Luz19 phage relative to the ancestral PAO1, with no significant difference in twitching motility measured between the positive-phage groups with/without SA (p > 0.05) or between non-phage conditions and the ancestral control p > 0.05). Boxplot ranges are 1.5X interquartile ranges.

### Growth assays and area under the curve

Our statistical analysis of growth asked two questions. First – did strains evolved under phage selection develop resistance. When compared to the ancestor, treatment with the ancestral phage resulted in significantly increased growth in lines evolved under phage pressure. (**Fig. 2**). Further, the addition or absence of phage produced no significant difference in growth in lines evolved with the phage. This is consistent with phage resistance. Our second question, did adaptation to the media or the presence of *SA* alter phage resistance? Our analysis revealed highly significant effects of both Treatment (F=132.73, df=3, p<2.22e-16) and Condition (F=41.83, df=4, p<2.22e-16) on the area under the growth curve, as well as significant Treatment x Condition interaction (F=37.66, df=12, p<2.22e-16). This indicates that not only do the individual treatments and co-evolution conditions affect growth outcomes, but the effect of treatment varies depending on the evolutionary history. (**Fig 2**). Subsequent post-hoc analysis shows that these results are due to phage resistance but only in lines which evolved under phage pressure. This increased resistance was observed in both the +Luz19 and +*SA*+Luz19 treatments compared to all phage-naïve lines, including ancestral PAO1 (p = 6.30e-4 and p = 1.65e-7, respectively), the PAO1t10 control (p = 1.23e-5 and p = 2.26e-9, respectively), and *PA* evolved with *SA* (p = 6.47e-5 and p = 1.35e-8, respectively). Importantly, the presence of *S. aureus* during evolution did not significantly alter phage resistance, our analysis found no significant difference between the +Luz19 and +*SA*+Luz19 conditions (p > 0.05), indicating that co-evolution with *S. aureus* did not alter resistance to the phage. Neither *SA* co-culture nor the media alone increased resistance to the phage relative to the ancestral PAO1 (p > 0.05). These results consistently demonstrate the enhanced phage resistance in phage positive lines, regardless of the presence of *SA* during evolution.

**Fig. 2.**
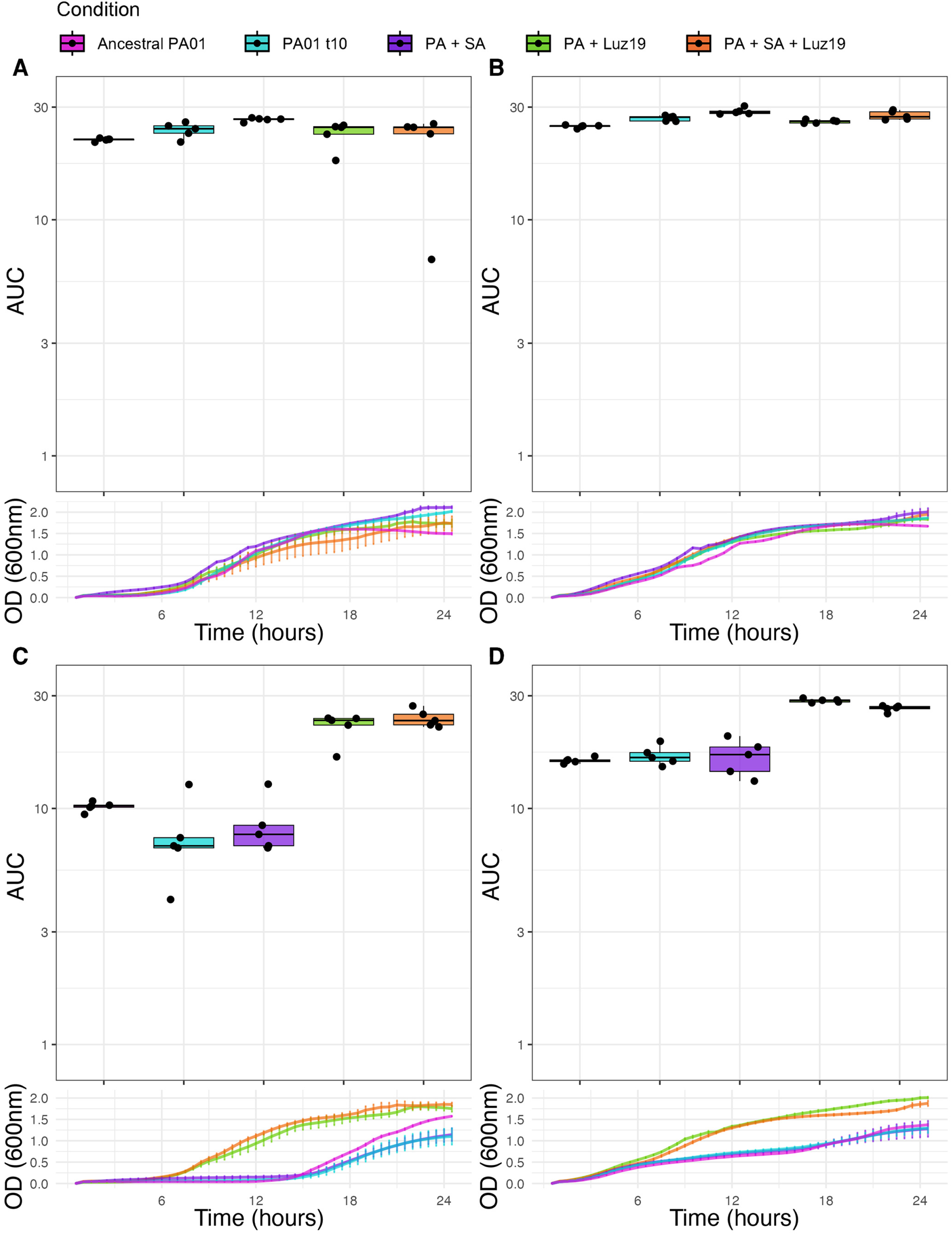
Growth assays and area under the curve (AUC). A.) Control treatment AUC boxplots (above) and mean growth curves (below). *PA* from the five conditions were grown in LB with no additional challenge – *PA* evolved alongside *SA* had significantly higher AUC than all other conditions (p < 0.05). **B.) *SA* treatment AUC boxplots (above) and mean growth curves (below)**. *PA* grown in LB with *SA* added. *PA* evolved with *SA* had significantly greater AUC than the ancestral (p < 0.05) but not significantly different from the transfer control PAO1t10 (p > 0.05).). **C.) Luz19 treatment AUC boxplots (above) and mean growth curves (below)**. Conditions subject to co-evolution with Luz19 (+Luz19 and +*SA+*Luz19) had significantly greater AUC than those not evolved with the phage (p < 0.05) and were not significantly different from one another (p > 0.05). There was no significant difference between the AUC of conditions not previously exposed to the phage (p > 0.05) **D.) *SA &* Luz19 treatment AUC boxplots (above) and mean growth curves (below)**. Conditions subject to co-evolution with Luz19 (+Luz19 and +*SA+*Luz19) had significantly greater AUC than those not evolved with the phage (p < 0.05) and were not significantly different from one another (p > 0.05). There was no significant difference between the AUC of conditions not previously exposed to the phage (p > 0.05). Boxplot ranges are the 1.5X interquartile ranges, error bars on the growth curves are Standard Error.

### Genomics

Sequencing revealed a clear signature of evolution in the presence of the phage (**Fig. 3**). Three SNP variations between two genes directly involved in type IV pilus biogenesis, *pilR* and *pilZ*, were defined in the conditions subject to co-evolution with the phage. In the two conditions with phage, 8 out of 10 replicates (5/5 +Phage, 3/5 +Phage/+*SA*) had SNPs in *pilR*, the response regulator element of the PilSR two component system, responsible for activation of *pilA* transcription [37].

**Fig. 3.**
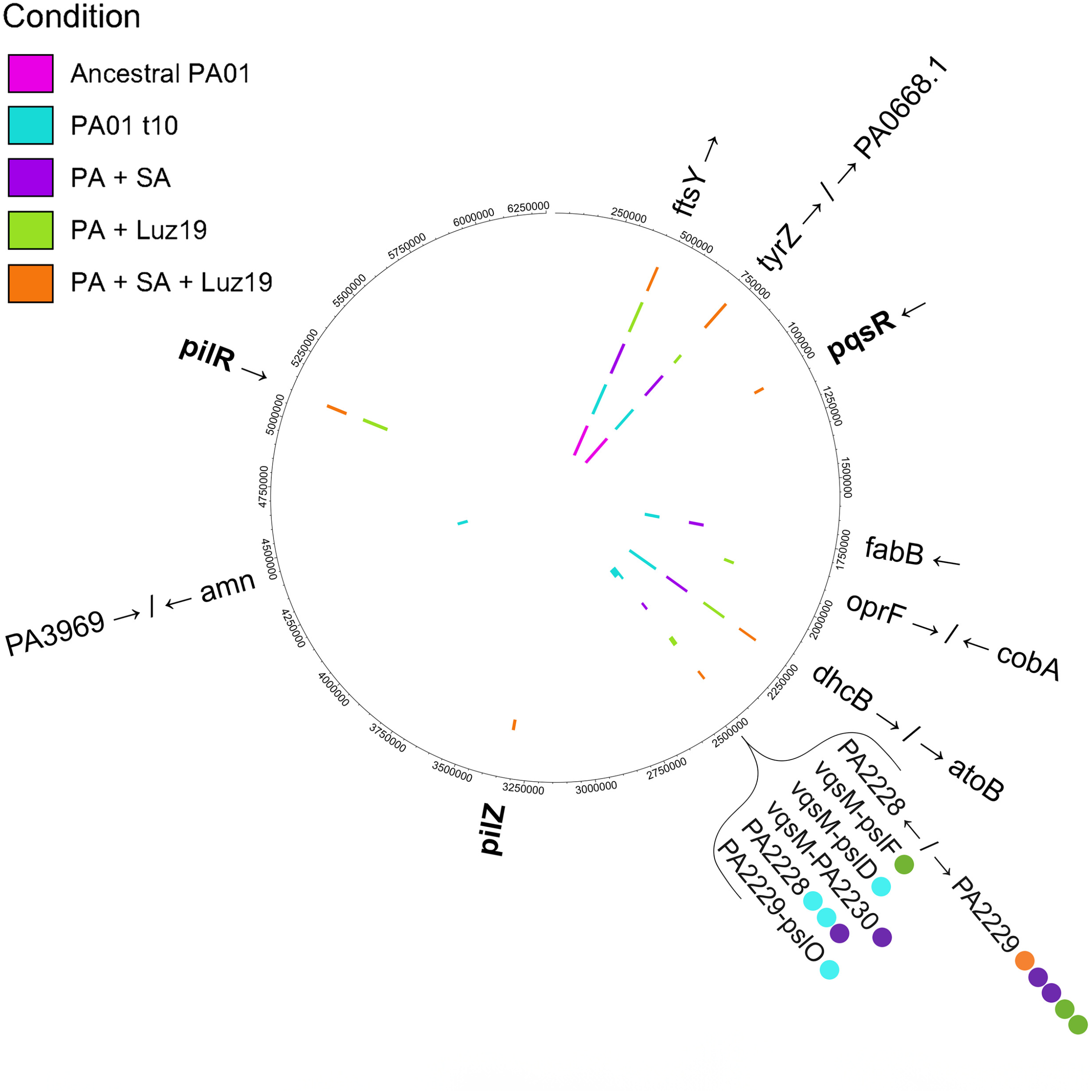
Chromosomal single nucleotide polymorphisms (SNPs) defined for all co-evolution condition replicates and the ancestral PAO1. SNPs were determined using breseq. ***pilR***. SNPs of two variations in 5 out of 5 (5/5) *+*Luz19 co-evo replicates and 3/5 *+SA+*Luz19 co-evo replicates. In all but one of these replicate sequences, the SNP in *pilR* is C192Y (TGC→TAC). Replicate *B* of the +Luz19 condition had a different SNP in *pilR*, P189Q (CCG→CAG). Of the replicates in the two conditions that were subject to co-evolution with Luz19, two lacked a SNP in *pilR*, both in the +*SA*+Luz19 condition. Replicate *B* instead had a 197 bp deletion in ***pilZ***. Replicate *A* of +*SA*+Luz19 was the only replicate co-evolved alongside Luz19 that sequencing did not reveal a *pil* gene mutation, instead, a PQS system mutation was identified in ***pqsR***.

Of the remaining two replicates, one (replicate A) had a 197 bp deletion in *pilZ*. The Δ197 bp *pilZ* replicate showed significantly attenuated twitching motility (p < 0.05) with no significant motility difference between it and the *pilR* mutants (**Fig. 3**).

Replicate B of the +*SA*+Luz19 condition lacked detectable SNPs in genes directly related to type IV pilus biogenesis, structure, or function. However, it did have a unique SNP in *pqsR*, which is a global virulence regulator involved in quorum sensing [38]. It is currently unclear what specific impact the *pqsR* mutation has on type IV pili expression or function, though the replicate did show roughly the same attenuation of twitching motility (**Fig. 1**) and resistant growth curves (**Fig. 2**)

## Discussion

Phage Steering is the use of phage to reduce or remove pathogenic behaviors from a bacterial population. Fundamentally, it works by phage making use of a bacterial product, typically an outer membrane expressed structure, to complete a full infection cycle. Bacteria are therefore under a selective pressure to modify the structure – engendering resistance to the phage. This modification also typically leads to a reduction in effectiveness of the structure in its evolved function. Should the structure not provide an actual fitness benefit in the environment in which the phage is applied there is no selective pressure to restore the phenotype. In infections like those seen in the Cystic Fibrosis lung, many virulence factors of *PA* do not provide a fitness benefit when considered in isolation, only contributing to nonadaptive virulence, increasing morbidity of people with CF [39]. This is likely due to *PA* not having evolved in the CF lung, as the CF lung itself is, in evolutionary terms, incredibly new. Before the 1930s, the majority of people with CF died before the lung became chronically infected with *PA* [40]. However, virulence factors such as type IV pili likely evolved to help bacteria deal with inter species competition [41-43]. Therefore, a central requirement of Phage Steering, that negative fitness effects are outweighed by the benefit of phage resistance, might not hold true if *PA* is in competition with other bacteria at the time of phage application.

To investigate how Phage Steering works during multi-species competition, we designed a simple experimental setup that provided the most favorable conditions for it to work. We conducted a co-evolutionary experiment involving either the phage Luz19 and bacterium *SA*, both in combination, alone or neither. The major result of our work is that regardless of the presence of *SA*, the phage still steered *PA* away from the expression of type IV pili (**Fig. 1**). We measured the phenotypic changes in type IV pili with twitching motility and confirmed phage resistance emerged as effectively in both treatments that had phage, indicating again that the presence of *SA* did not alter this outcome. Our growth analysis supports that phage resistance is similar with or without *SA* present (**Fig. 2**). We did see an increase in variability of phage resistance in the lines without phage, likely due to weak selection or drift (**Fig. 2C**). Our growth curves of isolates evolved without phage showed an increased AUC when tested in the presence of *SA* (**Fig. 2D**). This is due to the growth of the *SA*. Finally, we genomically characterized individual isolates after the evolution experiment and found the mutations selected by Luz19 were found in pili regulatory genes.

Examining these specific mutations in more detail, the gene product of *pilA*, is the major subunit, pilin (PilA) that comprises the type IV pilus shaft [44, 45]. PilS is located within the inner membrane of *PA* and is responsible for the detection of surplus pilin/PilA subunits localized to the inner membrane [37]. In the absence of surplus PilA, PilS autophosphorylates, and subsequently transfers the phosphate group to the response regulator, PilR [37]. PilR then binds to a specific sequence proximal to the promoter of *pilA;* and with the assistance of the alternative sigma factor, σ54, *pilA* is transcribed [37] (**Fig. 4**). Two unique SNPs in *pilR* were found in the phage-positive replicates, both point substitutions that result in amino acid changes. Of the eight *pilR* SNPs, seven were C192Y(TGC→TAC) with the sole outlier, P189Q (CCG→CAG). There was no significant difference between the attenuation of twitching motility between the two *pilR* mutation variants.

**Fig 4.**
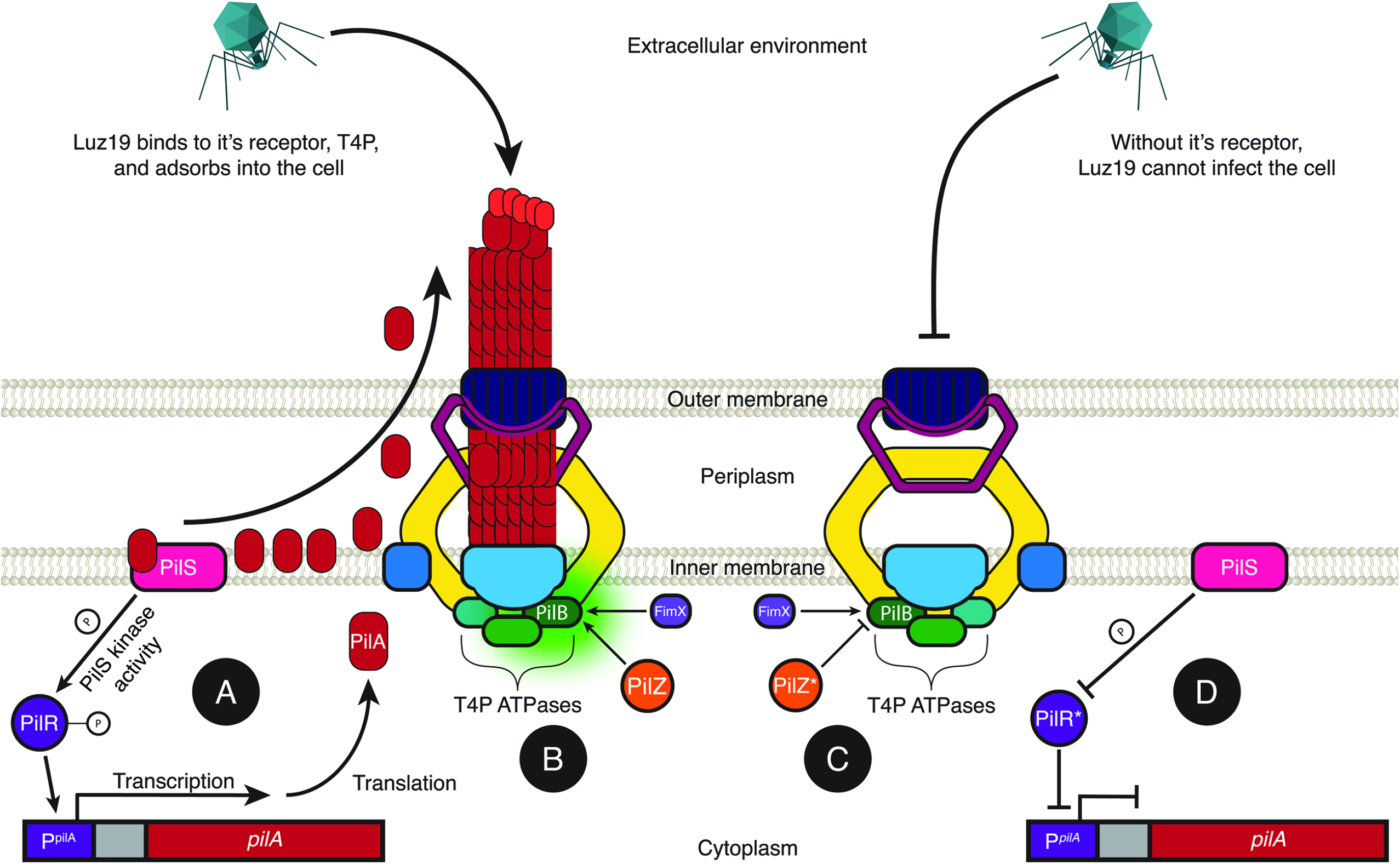
Mutations in key regulatory factors of T4P biogenesis confer phage resistance. A.) PilR serves as the response regulator of *pilA* transcription. The sensor kinase, PilS, phosphorylates PilR, activating it which in turn activate transcription of *pilA* which encodes for the major pilin subunit, PilA. **B.)** PilZ regulates T4P biogenesis. By binding to the extension ATPase, PilB, activates shaft polymerization. PilZ and FimX are necessary for PilB activation. **C.)** Without PilZ (PilZ*) binding, PilB remains inactive and energy is not generated for shaft assembly. **D.)** Defective PilR does not activate transcription of *pilA*, no PilA subunits are synthesized and thus the shaft is not assembled.

The gene product of *pilZ* is a critical regulatory element in the biogenesis of type IV pili [46]. PilZ interacts with PilB, an ATPase localized to the cytoplasmic face of the inner membrane that provides the energy necessary for transferring PilA subunits from the inner membrane to the base of the growing pilus polymer for pilus extension (**Fig. 4**) [46]. In the absence of functional PilZ, PilA subunits accumulate in the membrane but fail to assemble into a pilus shaft, resulting in loss of surface pili and associated phenotypes such as twitching motility and susceptibility to type IV pili-dependent phages [46] (**Fig. 4**). While the exact mechanism by which PilZ mediates PilB activity has yet to be fully elucidated, recent findings have shed significant light on the process. Hendrix et al. identified PlzR, a novel regulator, which binds to PilZ and modulates its function. Under elevated c-di-GMP levels, PlzR is transcriptionally induced and binds PilZ, preventing productive PilZ-PilB interaction, thereby inhibiting pilus assembly [47]. In contrast, at low c-di-GMP levels, FimX – another c-di-GMP-binding protein – interacts directly with PilB to promote pilus extension, suggesting that PilZ and FimX act as conditional regulators of the same extension ATPase [47]. Together, these findings establish PilZ as a central mediator linking secondary messenger signaling to pilus biogenesis. This layered regulatory model, proposed by Hendrix et al, is supported by our finding of a 197 bp deletion in PilZ in one of the Luz19-challenged evolutionary lineages (replicate A of +*SA*+Luz19). This loss-of-function mutation conferred phage resistance and led to significantly attenuated twitching motility. The emergence of this deletion under strong phage selection highlights PilZ’s essentiality for type IV pilus biogenesis and thus its disruption serves as a rapid path to type IV pili-dependent phage resistance.

Slightly unexpected is the appearance of a quorum sensing mutation in the *pqsR* regulator, which is also known as *mvfR*, a regulator in the Pseudomonas Quinolone Signal (PQS) quorum sensing system. The phage pf, also a type IV pilus binding phage has been shown to generate mutations within the PQS system [48], therefore there is likely some interaction between the PQS system and type IV pili. Guo et al. showed that while adding exogenous PQS did not alter the twitching motility, *pqsR* mutants had attenuated twitching [49]. It is somewhat surprising that this mutation was seen in the *SA*-positive line. PQS is a pivotal quorum sensing molecule in *PA* that regulates biofilm formation along with a plethora of virulence factors [50]. The role that PQS plays in the killing of *SA* is complex with many PQS regulated factors contributing to *PA*’s competitive advantage. In brief, PQS has antimicrobial properties against *SA* via the action of 2-heptyl-4-hydroxyquinoline (HHQ) and 2-heptyl-4-hydroxyquinoline N-oxide (HQNO) [51-56]. In PQS mutants (Δ*pqsA* and Δ*pqsH*), a reduced ability to kill *SA* has been shown relative to WT *PA* strains [38, 52]. While PQS plays a critical role in *PA*’s ability to kill *SA*, it is part of a complex network of QS molecules and virulence factors influenced by the PQS quorum sensing system. Notably, the antimicrobial activity of PQS and its derivatives is modulated by environmental factors, particularly iron availability [54]. Consequently, competition studies and further investigations using media that more accurately mimic *in vivo* nutrient-limited environments, such as SCFM2, may illuminate potential evolutionary trade-offs or reveal vulnerabilities in Luz19-resistant *PA* harboring *pqsR* mutations.

These phenotypic and genomic data point us towards the conclusion that Phage Steering is effective in the presence of *SA* when targeting the type IV pilus. In future work we will expand this to include other bacterial pathogens and commensals, as well as working in media which better mimics infection sites, such as SCFM2 [39]. We know that the essential genome of bacteria, as measured by TN-seq, is dependent on environment, both the biotic (interspecies competition) and the abiotic [57]. It is possible that infection environments which include competition from pathogens and commensals, a host immune system, and nutritional limitations might impose a fitness cost on the loss of structures which phages can target.

## References

1. Betts, A., O. Kaltz, and M.E. Hochberg, Contrasted coevolutionary dynamics between a bacterial pathogen and its bacteriophages. Proc Natl Acad Sci U S A, 2014. 111(30): p. 11109–14.

2. Rohde, C., et al., Expert Opinion on Three Phage Therapy Related Topics: Bacterial Phage Resistance, Phage Training and Prophages in Bacterial Production Strains. Viruses, 2018. 10(4).

3. Medel, R., et al., Arms Race Coevolution: The Local and Geographical Structure of a Host–Parasite Interaction. Evolution: Education and Outreach, 2010. 3(1): p. 26–31.

4. Thompson, J.N., The geographic mosaic of coevolution. 2019: University of Chicago Press.

5. Thompson, J.N. and M. Pagel, Coevolution. Encyclopedia of Evolution. Oxford University Press, Oxford, 2002.

6. Chao, L., B.R. Levin, and F.M. Stewart, A Complex Community in a Simple Habitat: An Experimental Study with Bacteria and Phage. Ecology, 1977. 58(2): p. 369–378.

7. Horne, M.T., Coevolution of Escherichia coli and bacteriophages in chemostat culture. Science, 1970. 168(3934): p. 992–3.

8. Koskella, B. and M.A. Brockhurst, Bacteria-phage coevolution as a driver of ecological and evolutionary processes in microbial communities. FEMS Microbiol Rev, 2014. 38(5): p. 916–31.

9. Lenski, R.E. and B.R. Levin, Constraints on the Coevolution of Bacteria and Virulent Phage: A Model, Some Experiments, and Predictions for Natural Communities. The American Naturalist, 1985. 125(4): p. 585–602.

10. Schrag, S.J. and J.E. Mittler, Host-parasite coexistence: the role of spatial refuges in stabilizing bacteria-phage interactions. The American Naturalist, 1996. 148(2): p. 348–377.

11. Spanakis, E. and M.T. Horne, Co-adaptation of Escherichia coli and coliphage lambda vir in continuous culture. J Gen Microbiol, 1987. 133(2): p. 353–60.

12. Hall, A.R., et al., Host-parasite coevolutionary arms races give way to fluctuating selection. Ecol Lett, 2011. 14(7): p. 635–42.

13. Hall, Alex R., Pauline D. Scanlan, and A. Buckling, Bacteria‐Phage Coevolution and the Emergence of Generalist Pathogens. The American Naturalist, 2011. 177(1): p. 44–53.

14. Hall, A.R., et al., Host-parasite coevolutionary arms races give way to fluctuating selection. Ecology Letters, 2011. 14(7): p. 635–642.

15. Gurney, J., et al., Steering Phages to Combat Bacterial Pathogens. Trends Microbiol, 2020. 28(2): p. 85–94.

16. Gurney, J., et al., Phage steering of antibiotic-resistance evolution in the bacterial pathogen, Pseudomonas aeruginosa. Evol Med Public Health, 2020. 2020(1): p. 148–157.

17. Diggle, S.P. and M. Whiteley, Microbe Profile: Pseudomonas aeruginosa: opportunistic pathogen and lab rat. Microbiology (Reading), 2020. 166(1): p. 30–33.

18. DeLeon, S., et al., Synergistic interactions of Pseudomonas aeruginosa and Staphylococcus aureus in an in vitro wound model. Infect Immun, 2014. 82(11): p. 4718–28.

19. Hatziagorou, E., et al., Changing epidemiology of the respiratory bacteriology of patients with cystic fibrosis-data from the European cystic fibrosis society patient registry. J Cyst Fibros, 2020. 19(3): p. 376–383.

20. Dalton, T., et al., An in vivo polymicrobial biofilm wound infection model to study interspecies interactions. PLoS One, 2011. 6(11): p. e27317.

21. Korgaonkar, A., et al., Community surveillance enhances Pseudomonas aeruginosa virulence during polymicrobial infection. Proc Natl Acad Sci U S A, 2013. 110(3): p. 1059–64.

22. Korgaonkar, A.K. and M. Whiteley, Pseudomonas aeruginosa enhances production of an antimicrobial in response to N-acetylglucosamine and peptidoglycan. J Bacteriol, 2011. 193(4): p. 909–17.

23. Sibley, C.D., et al., Discerning the complexity of community interactions using a Drosophila model of polymicrobial infections. PLoS Pathog, 2008. 4(10): p. e1000184.

24. Alseth, E.O., et al., The impact of phage and phage resistance on microbial community dynamics. PLOS Biology, 2024. 22(4): p. e3002346.

25. Blazanin, M. and P.E. Turner, Community context matters for bacteria-phage ecology and evolution. ISME J, 2021. 15(11): p. 3119–3128.

26. Langendonk, R.F., D.R. Neill, and J.L. Fothergill, The Building Blocks of Antimicrobial Resistance in Pseudomonas aeruginosa: Implications for Current Resistance-Breaking Therapies. Front Cell Infect Microbiol, 2021. 11: p. 665759.

27. Fernandez-Billon, M., et al., Mechanisms of antibiotic resistance in Pseudomonas aeruginosa biofilms. Biofilm, 2023. 5: p. 100129.

28. Grosso-Becerra, M.V., et al., Pseudomonas aeruginosa clinical and environmental isolates constitute a single population with high phenotypic diversity. BMC Genomics, 2014. 15: p. 318.

29. Qin, S., et al., Pseudomonas aeruginosa: pathogenesis, virulence factors, antibiotic resistance, interaction with host, technology advances and emerging therapeutics. Signal Transduct Target Ther, 2022. 7(1): p. 199.

30. Chibeu, A., et al., The adsorption of Pseudomonas aeruginosa bacteriophage phiKMV is dependent on expression regulation of type IV pili genes. FEMS Microbiol Lett, 2009. 296(2): p. 210–8.

31. Lammens, E., et al., Representational Difference Analysis (RDA) of bacteriophage genomes. J Microbiol Methods, 2009. 77(2): p. 207–13.

32. Alm, R.A. and J.S. Mattick, Identification of two genes with prepilin-like leader sequences involved in type 4 fimbrial biogenesis in Pseudomonas aeruginosa. J Bacteriol, 1996. 178(13): p. 3809–17.

33. Weisberg, J.F.a.S., An {R} Companion to Applied Regression. Third ed. 2019: Sage.

34. Sprouffske, K. and A. Wagner, Growthcurver: an R package for obtaining interpretable metrics from microbial growth curves. BMC Bioinformatics, 2016. 17: p. 172.

35. Deatherage, D.E. and J.E. Barrick, Identification of mutations in laboratory-evolved microbes from next-generation sequencing data using breseq. Methods Mol Biol, 2014. 1151: p. 165–88.

36. Stover, C.K., et al., Complete genome sequence of Pseudomonas aeruginosa PAO1, an opportunistic pathogen. Nature, 2000. 406(6799): p. 959–64.

37. Boyd, J.M. and S. Lory, Dual function of PilS during transcriptional activation of the Pseudomonas aeruginosa pilin subunit gene. J Bacteriol, 1996. 178(3): p. 831–9.

38. Lin, J., et al., The Pseudomonas Quinolone Signal (PQS): Not Just for Quorum Sensing Anymore. Front Cell Infect Microbiol, 2018. 8: p. 230.

39. Turner, K.H., et al., Essential genome of Pseudomonas aeruginosa in cystic fibrosis sputum. Proc Natl Acad Sci U S A, 2015. 112(13): p. 4110–5.

40. Davis, P.B., Cystic Fibrosis Since 1938. American Journal of Respiratory and Critical Care Medicine, 2006. 173(5): p. 475–482.

41. Yarrington, K.D., T.N. Shendruk, and D.H. Limoli, The type IV pilus chemoreceptor PilJ controls chemotaxis of one bacterial species towards another. PLoS Biol, 2024. 22(2): p. e3002488.

42. Limoli, D.H., et al., Interspecies interactions induce exploratory motility in Pseudomonas aeruginosa. Elife, 2019. 8.

43. Sanchez-Pena, A., et al., Pseudomonas aeruginosa surface motility and invasion into competing communities enhance interspecies antagonism. mBio, 2024. 15(9): p. e0095624.

44. Sastry, P.A., et al., Comparative studies of the amino acid and nucleotide sequences of pilin derived from Pseudomonas aeruginosa PAK and PAO. J Bacteriol, 1985. 164(2): p. 571–7.

45. Strom, M.S. and S. Lory, Mapping of export signals of Pseudomonas aeruginosa pilin with alkaline phosphatase fusions. J Bacteriol, 1987. 169(7): p. 3181–8.

46. Guzzo, C.R., et al., PILZ Protein Structure and Interactions with PILB and the FIMX EAL Domain: Implications for Control of Type IV Pilus Biogenesis. Journal of Molecular Biology, 2009. 393(4): p. 848–866.

47. Hendrix, H., et al., PlzR regulates type IV pili assembly in Pseudomonas aeruginosa via PilZ binding. Nat Commun, 2024. 15(1): p. 8717.

48. Schwartzkopf, C.M., et al., Inhibition of PQS signaling by the Pf bacteriophage protein PfsE enhances viral replication in Pseudomonas aeruginosa. Mol Microbiol, 2024. 121(1): p. 116–128.

49. Guo, Q., et al., PqsR-dependent and PqsR-independent regulation of motility and biofilm formation by PQS in Pseudomonas aeruginosa PAO1. Journal of Basic Microbiology, 2014. 54(7): p. 633–643.

50. Pesci, E.C., et al., Quinolone signaling in the cell-to-cell communication system of Pseudomonas aeruginosa. Proc Natl Acad Sci U S A, 1999. 96(20): p. 11229–34.

51. Filkins, L.M., et al., Coculture of Staphylococcus aureus with Pseudomonas aeruginosa Drives S. aureus towards Fermentative Metabolism and Reduced Viability in a Cystic Fibrosis Model. J Bacteriol, 2015. 197(14): p. 2252–64.

52. Haussler, S. and T. Becker, The pseudomonas quinolone signal (PQS) balances life and death in Pseudomonas aeruginosa populations. PLoS Pathog, 2008. 4(9): p. e1000166.

53. Nairz, M., et al., The struggle for iron - a metal at the host-pathogen interface. Cell Microbiol, 2010. 12(12): p. 1691–702.

54. Nguyen, A.T., et al., Cystic Fibrosis Isolates of Pseudomonas aeruginosa Retain Iron-Regulated Antimicrobial Activity against Staphylococcus aureus through the Action of Multiple Alkylquinolones. Front Microbiol, 2016. 7: p. 1171.

55. Nguyen, A.T. and A.G. Oglesby-Sherrouse, Interactions between Pseudomonas aeruginosa and Staphylococcus aureus during co-cultivations and polymicrobial infections. Appl Microbiol Biotechnol, 2016. 100(14): p. 6141–6148.

56. Otto, B.R., A.M. Verweij-van Vught, and D.M. MacLaren, Transferrins and heme-compounds as iron sources for pathogenic bacteria. Crit Rev Microbiol, 1992. 18(3): p. 217–33.

57. Ibberson, C.B., et al., Co-infecting microorganisms dramatically alter pathogen gene essentiality during polymicrobial infection. Nat Microbiol, 2017. 2: p. 17079.

