## Supplemental information for "Phage Steering in the Presence of a Competing Bacterial Pathogen"

| **Oligo name** | **Sequence (5’-3’)** |
| --- | --- |
| Luz7 17892-17913 FWD 1 | gcccgtgctctgtacgttgctt |
| Luz7 18044-18065 REV 1 | gcagacgcagcagctggttgta |

SI table 1. Primers used in qPCR quantification of Phage concentration.

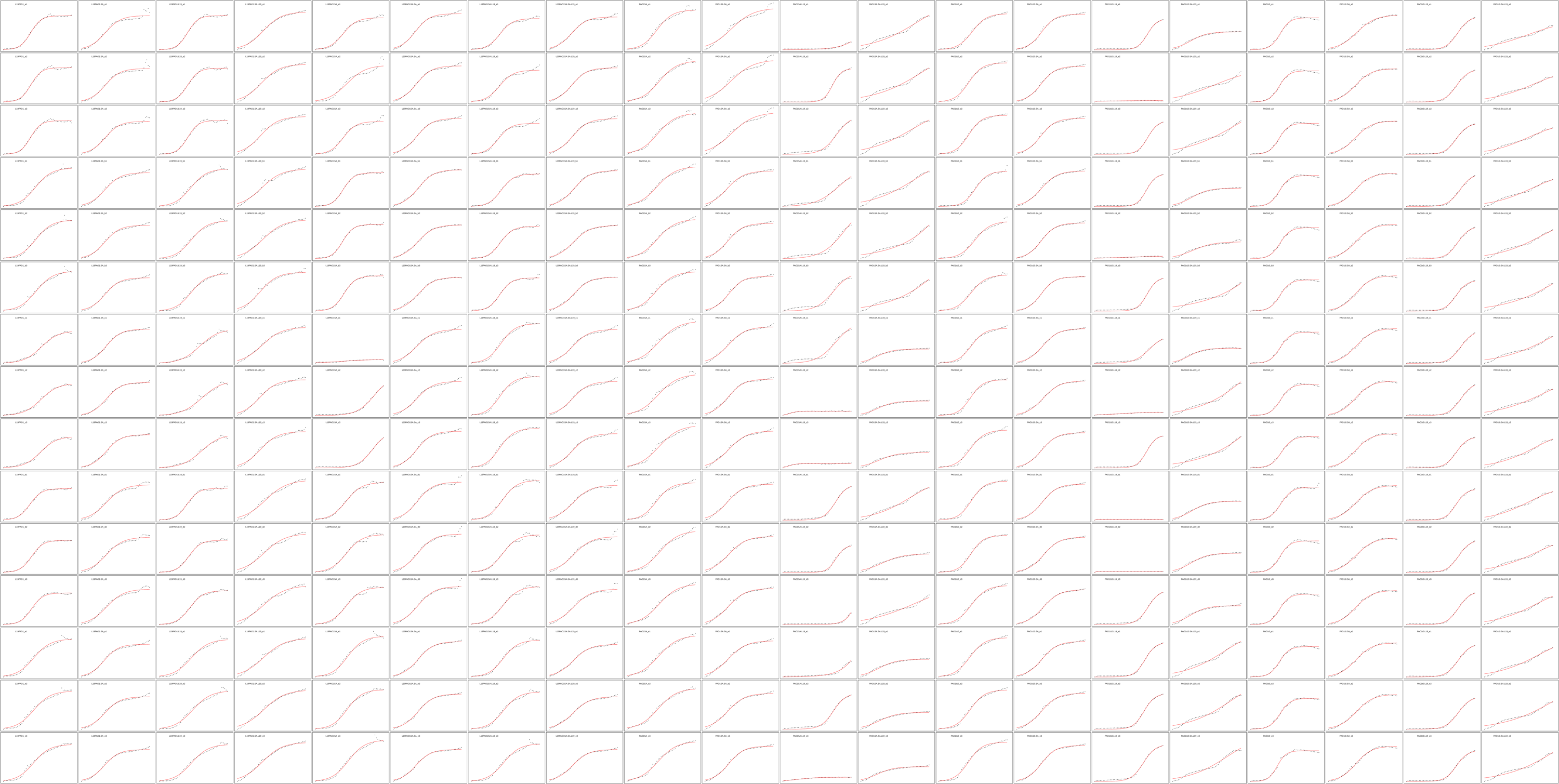

**SI Fig 1. Growthcurver outputs of all Condition x Treatment combination with each line assayed in triplicate. Plots show growthcurver logistic curve modeling (red) using individual OD_600_ measurements taken every 30 minutes for 24.5 hours (black dots). Each column represents a different combination of Condition x Treatment with 5 lines each tested in triplicate for a total of 15 replicates per combination – not accounting for the pseudo-replication created by the assay triplicates.**

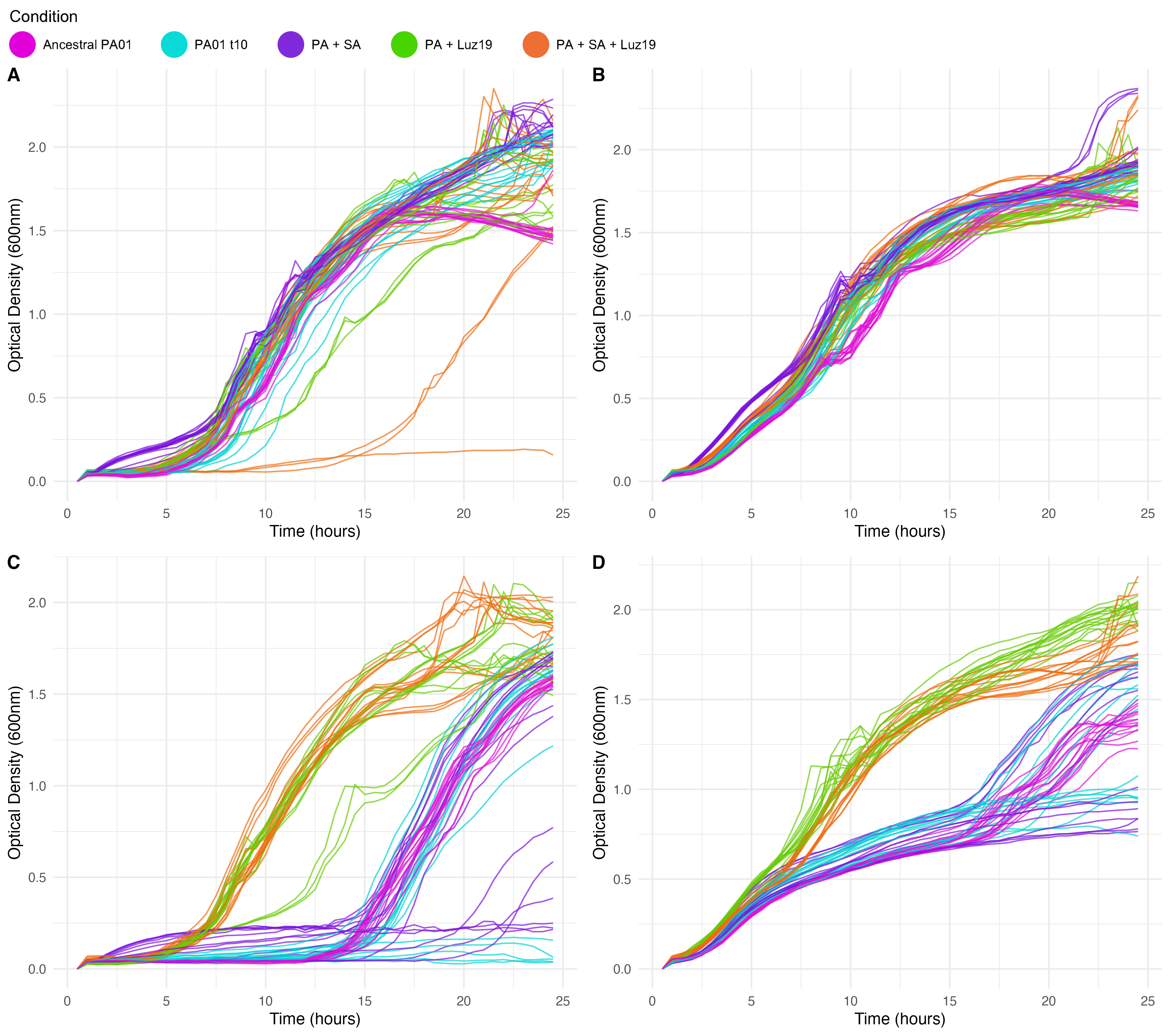

**SI Fig 2. Growth curves of each individual co-evolution condition line and their respective individual triplicates subject to each of the four treatments. A.)No Treatment. Curves are consistent with a few replicates of +*SA+*Luz19 exhibiting delayed growth. B.) *S. aureus* treatment. The absence of a nominal *PA* lag phase is notable, this is due to *SA* growth and absorbance during *PA* lag phase. No outliers of note. C.) Luz19 treatment. PA previously evolved with Luz19 exhibit growth curves resembling that seen in A. Replicates not previously exposed to the phage display delayed growth as phage resistant clones sweep through the population at the 15-20 hour mark or no growth is observed. Bottom right. *S. aureus* & Luz19 treatment. A lack of lag phase is once again notable as the *SA* dominates hours 0-5. In phage-evolved conditions, *PA* nominal curve takes over around the 5 hours mark. In contrast, phage-naïve conditions exhibit delayed growth – once again sweeps either take off around the 15-20 hour mark and take over or *PA* growth is not observed and *SA* continues to dominate.**

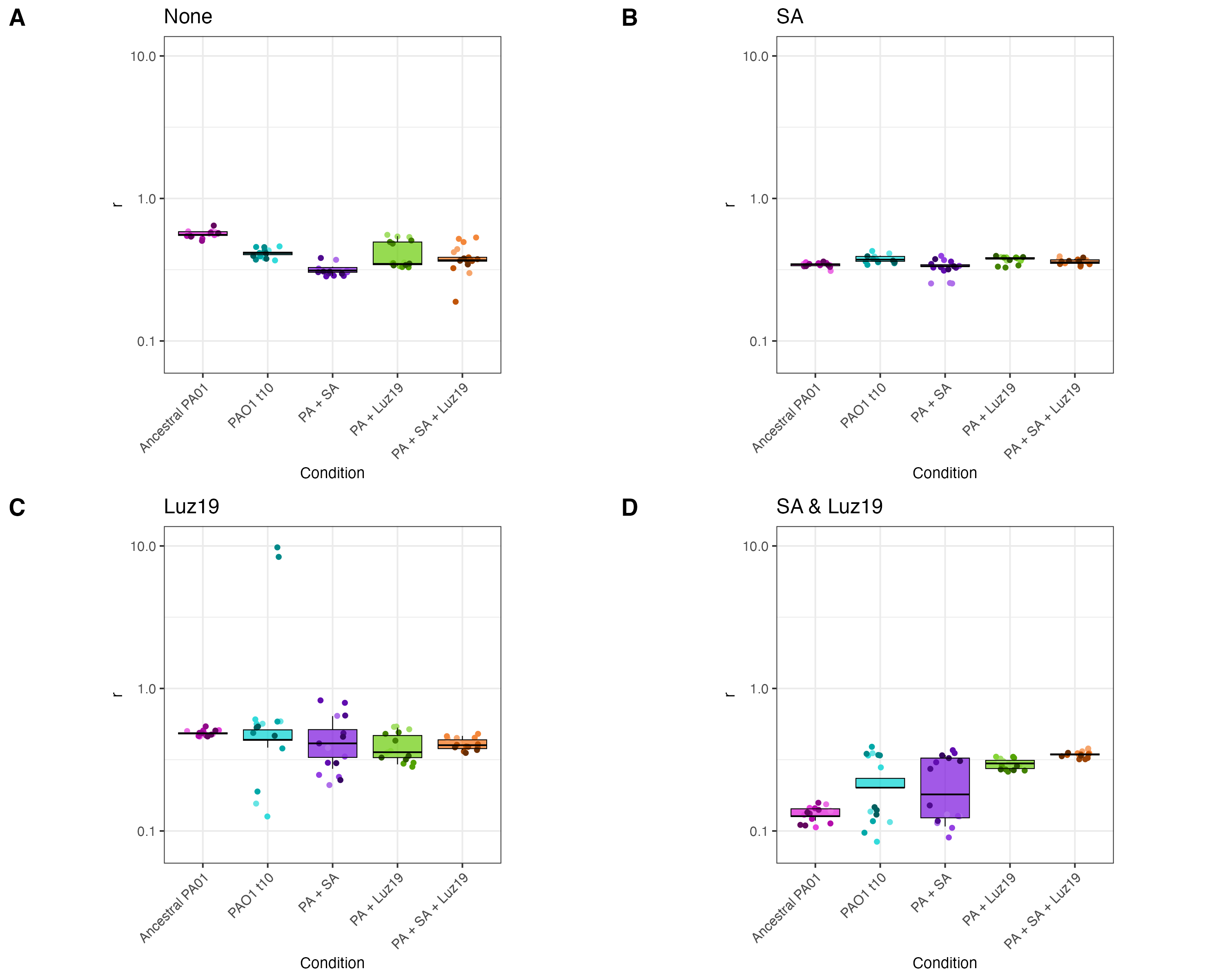

**SI Fig 3. Mean intrinsic growth rate of each coevolution condition line subject to the assay treatments plotted as boxplots with a logarithmic scaled Y-axis.** **A.) No Treatment. B.) *S. aureus* treatment. C.) Luz19 treatment. With the Luz19 treatment added, two replicates within the transfer control condition PAO1t10 produce outliers that skew the data. These outliers occur due to phage resistant clones sweeping through the population late causing a delayed log phase that is predicted by Growthcurver to continue exponential growth far beyond the real carrying capacity. D.) *S. aureus* & Luz19 treatment.**

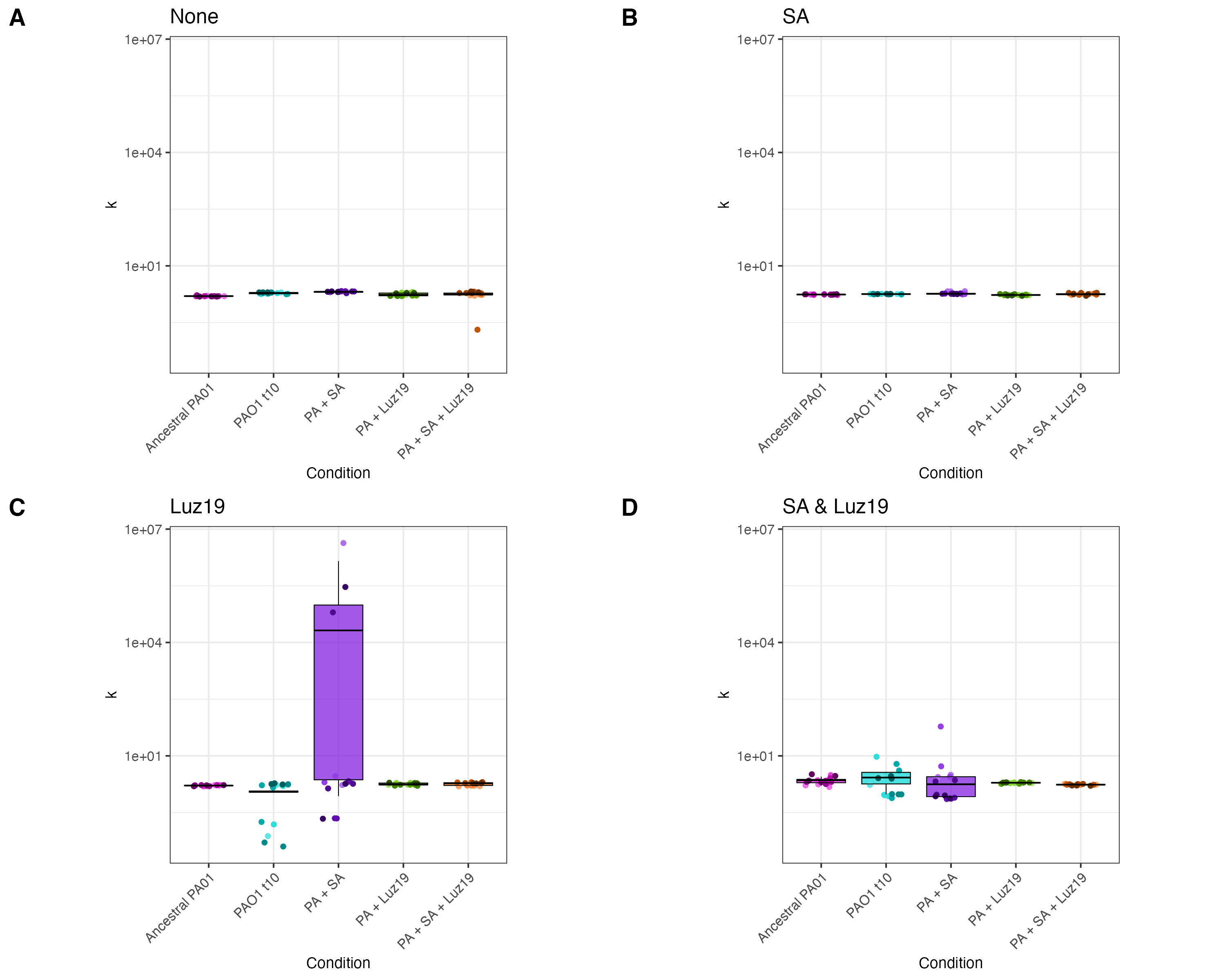

**SI Fig 4. Boxplots with linear Y axis of Growthcurver predicted carrying capacity (k) of each condition subject to the four treatments. A.)No Treatment. All conditions achieved a similar carrying capacity with no additional treatment. B.) *S. aureus* treatment. C.) Luz19 treatment. PA previously evolved with SA saw significantly greater predicted carrying capacity than the ancestor (PAO1t0) and PAo1t10 (p < 0.05). Bottom right. *S. aureus* & Luz19 treatment.**
